# Image-based pooled whole genome CRISPR screening for Parkin and TFEB subcellular localization

**DOI:** 10.1101/2020.07.02.184390

**Authors:** Gil Kanfer, Shireen A. Sarraf, Yaakov Maman, Heather Baldwin, Kory R. Johnson, Michael E. Ward, Martin Kampmann, Jennifer Lippincott-Schwartz, Richard J. Youle

## Abstract

Genome-wide CRISPR screens have transformed our ability to systematically interrogate human gene function, but are currently limited to a subset of cellular phenotypes. We report a novel pooled screening approach for a wider range of cellular and subtle subcellular phenotypes. Machine learning and convolutional neural network models are trained on the subcellular phenotype to be queried. Genome-wide screening then utilizes cells stably expressing dCas9 (CRISPRi), photoactivatable fluorescent protein (PA-mCherry), and a lentiviral guide RNA (gRNA) pool. Cells are screened by microscopy and classified by artificial intelligence (AI) algorithms, which precisely identify the genetically altered phenotype. Cells with the phenotype of interest are photoactivated, isolated via flow cytometry, and the gRNAs are identified by sequencing. A proof-of-concept screen accurately identified PINK1 as essential for Parkin recruitment to mitochondria. A genome-wide screen identified factors mediating TFEB relocation from the nucleus to the cytosol upon prolonged starvation. Twenty of the sixty-four hits called by the neural network model were independently validated, revealing new effectors of TFEB subcellular localization. This approach, AI-Photoswitchable Screening (AI-PS) offers a novel screening platform capable of classifying a broad range of mammalian subcellular morphologies, an approach largely unattainable with current methodologies at genome-wide scale.

## Introduction

Recent advances have expanded traditional genetic screens from bacteria and yeast to mammalian cells. RNAi, CRISPRi and CRISPR screens rely on two main strategies: arrayed and pooled screens. Arrayed screens are highly specific but require the production and, by definition, individual assortment of each RNAi or CRISPR guide separately, requiring high-throughput equipment not readily available to academic labs. Pooled screens are more facile but restricted to phenotypes that affect cell growth rates or viability or result in a fluorescence increase that allows isolation of hits from the population using FACS. Single-cell RNA based pooled screens are also useful to link genetic profiles to perturbations (Horlbeck et al., 2018; Datlinger et al., 2017; Dixit et al., 2016; Adamson et al., 2016). Recent reports describe new strategies that distinguish microscopic differences in images from pooled screens. Microfluidic separation of cells from the population (Ota et al., 2018; Nitta et al., 2018), *in situ* sequencing of guides in individual cells (Wang et al., 2019; Feldman et al., 2018; Emanuel et al., 2017), microscale cell carrier technology (Wheeler et al., 2020), link genotypes to images of cell phenotypes. Though elegant, these approaches have technical limitations and are not applicable at the whole genome scale.

Recent advances in machine learning, and particularly in deep learning (convolutional neural networks) (Caicedo et al., 2018; Bzdok et al., 2018) offer novel strategies for identifying individual cells with altered organelle morphology or subcellular protein localization. We developed a unique screening method to identify genetic perturbations of subcellular morphologies that is widely applicable and high-throughput. The method is divided into four steps: first, a morphology classification model is trained on single-cell images. Second, pools of CRISPRi-perturbed target cells are imaged sequentially, and the phenotypically selected cells are labeled by laser photoactivation of a fluorescent protein. Third, the photoactivated cells are sorted and fourth, the guides within phenotypically identified cells are amplified and sequenced. The decision to select cells is made on-the-fly by pre-trained classification models allowing screening of 1×10^6^ cells within 12 hours and the whole human genome in a week.

## RESULTS

### Building the single-cell imaging screening approach

We developed a new platform which assesses images of cells and uses machine learning to distinguish their subcellular phenotypes. Using laser activation of a fluorescent probe to denote the selected cell phenotypes and FACS to separate the cells for guide sequencing, one essentially converts the individual cells exposed to pooled CRISPRi libraries into an arrayed screen (Fig. 1). Using this approach, every imaged cell is referred to as an independent entity, and a predicted phenotype score is produced based on a classification machine-learning model (Fig. 1 e). Making the AI platform entails three steps: training and creation of the phenotype classification model; model deployment on pooled imaged cells; and validation of the model’s screening performance. We utilized Pink1-dependent Parkin translocation to mitochondria as a proof-of-concept (Fig. 1 b). In cells with unimpaired polarized mitochondria, Parkin is in the cytoplasm; however, upon mitochondrial depolarization, it translocates to mitochondria(Narendra et al., 2008) (Fig. 1 b,c). This binary switch in Parkin location is suitable for detection by a support vector machine (SVM) classification model. An SVM classification model was trained on images of cells with either cytosolic or mitochondrial GFP-Parkin. For each single-cell image, the program calculates a broad set of features based on measurements of the GFP signal pixels (Fig. 1 d,e). Feature variations were assessed, and redundant contributions were excluded from the model (Fig. S1a,b). Manually selected cell images annotated by phenotype and image features were computationally applied on a radial basis kernel SVM to create the classification model (Fig. S1 a and b). For this step, we generated an easy-to-use graphical user interface program to facilitate image segmentation, measurement and model building (Fig 2). The R-based script for image segmentation and analysis, and the SVM-classification model (Fig. 2b) were deployed on-the-fly to identify cells exhibiting the desired phenotype (GFP-Parkin mitochondrial localization). During live cell image acquisition, single cell images were captured following segmentation and stored on a local computer. The SVM-based model classified the individual cells and generated a mask corresponding to the live image field identifying the location of cells with the phenotype of interest (Fig. 1g, Extended Data Fig. 2a,b, and Video 1). In cells identified with this mask, PA-mCh was then laser photoactivated. This 10 second process was iterated across serial images of the entire chamber slide (an average of 500,000 cells for one subgenomic CRISPRi guide pool (Gilbert et al., 2014; Horlbeck et al., 2016)). Finally, the photoactivated samples were sorted using flow cytometry and deep sequenced to determine sgRNA abundance in the activated sample compared to untreated cells.

**Figure 1.**
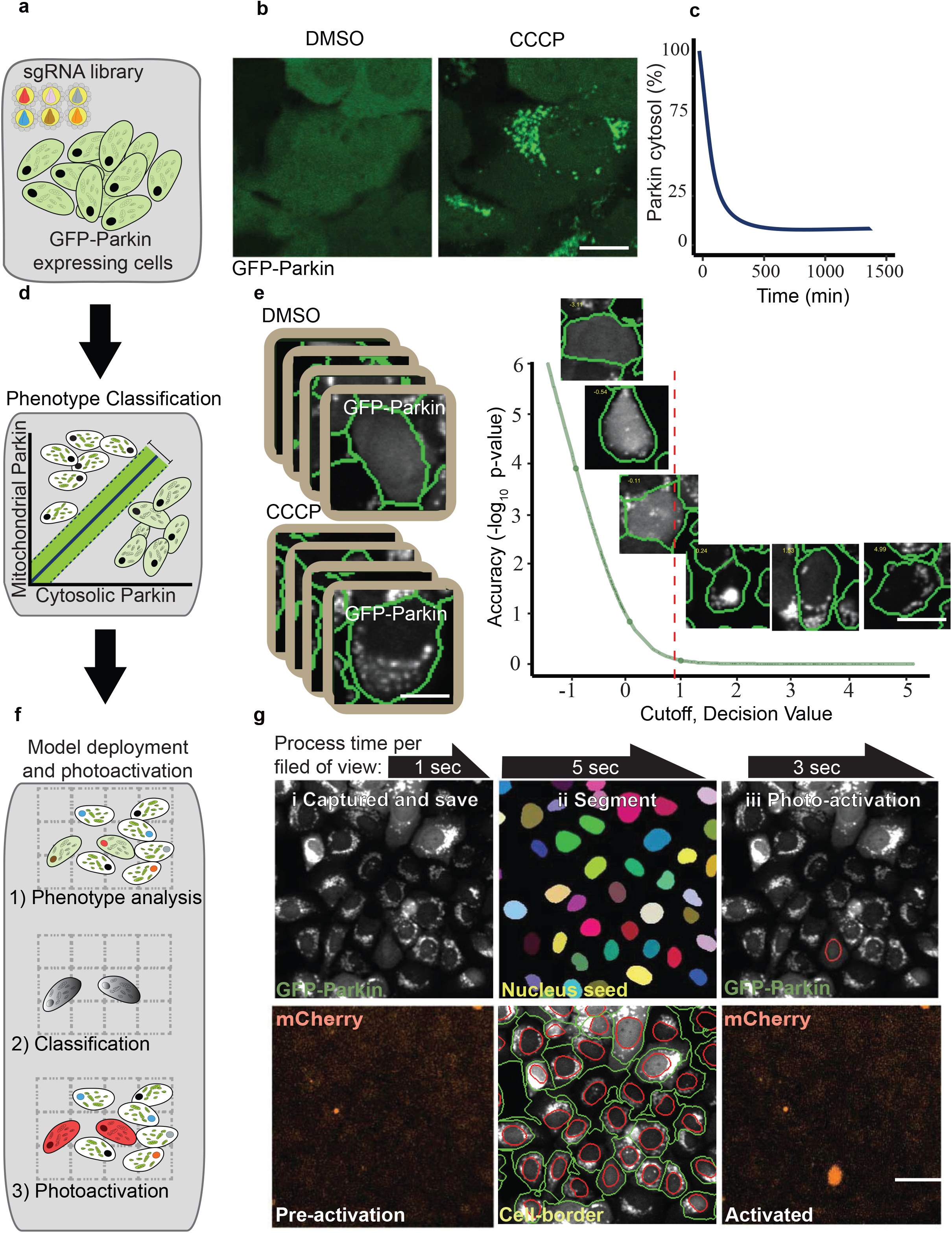
Machine learning genetic screening platform for Parkin localization - proof of principle screen. **(a)** Pooled sgRNAs infecting target cells. **(b)** GFP-Parkin U2OS cells treated with DMSO or CCCP (2 hr). Scale bar, 5 μm. **(c)** Percent of cytosolic Parkin over time after CCCP 10 μM treatment supplemented with 100 nM Bafilomycin A. **(d, e)** Single-cell image examples for the training set of phenotypes used for SVM classification. Cutoff determined by classification model accuracy (y axis). Cutoff used in this screen is a prediction score of 0.8 (also known as decision value), scale bar, 5 μm **(f, g)** Representative Field Of View (FOV) of GFP-Parkin U2OS cell screening procedure. i, Images were captured and saved on local computer. ii, Cells borders were identified (green circle surrounding cell border, red circle nucleus) following Nucleus segmentation (color labeled nuclei). SVM classification model was deployed and masked (red circle). iii, photoactivation of the SMV identified cell; scale bar, 10 μm

**Figure. 2.**
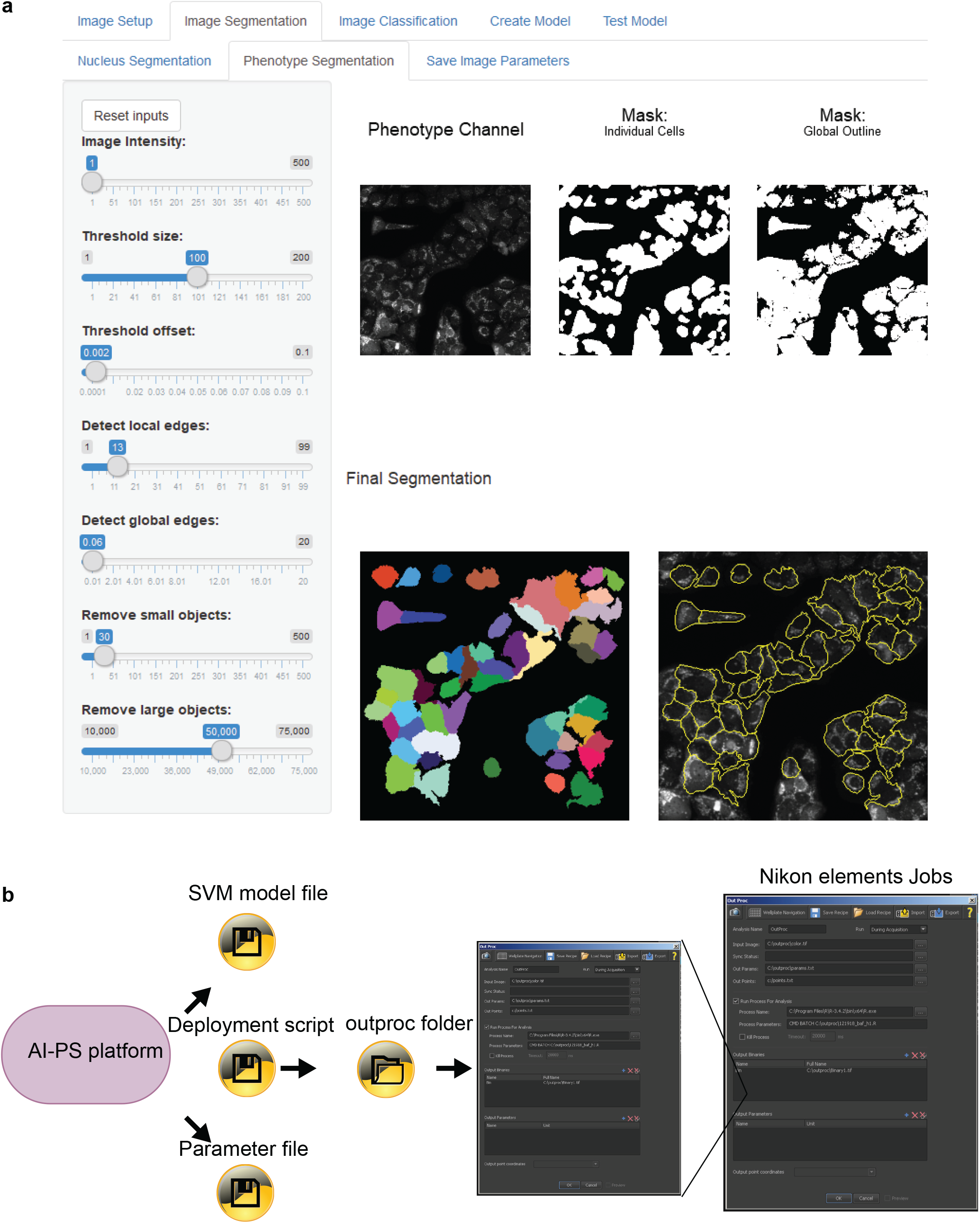
Shiny AI-PS Application and output files. **(a)** AI-PS GUI interface example. https://hab-gk-app.shinyapps.io/gk_shiny_app/ **(b)** Three deployment output files integrated with Nikon elements software.

### Parkin translocation screen validation

For platform validation, U2OS cells stably expressing GFP-Parkin, PA-mCh, and dCAS9 were infected with subpool of the version 2 CRISPRi library comprising 12,775 guides targeting kinases, phosphatases and the druggable genome (Horlbeck et al., 2016). Cells were treated with carbonyl cyanide m-chlorophenyl hydrazone (CCCP) to depolarize mitochondria and GFP-Parkin localization was assessed using the SVM classification model (Fig. 3 a). From approximately 200,000 cells, 1,132 were called, photoactivated, sorted and sequenced (Fig. 3 b). The most abundant sgRNAs identified in the photoactivated samples were targeted against PINK1 (Fig. 3 c), known to be required for Parkin translocation^16^, exhibiting a nearly 30-fold increase compared to the unsorted control sample (FDR adjusted p-value < 0.0001, Table S1). Thus, the single known Parkin modifier targeted by the subpool library, PINK1, was identified, validating the method. In addition, sample size estimation indicated that three biological repeats are sufficient for detecting the desired genetic link in our experimental setup (Fig. S2 d).

**Figure 3.**
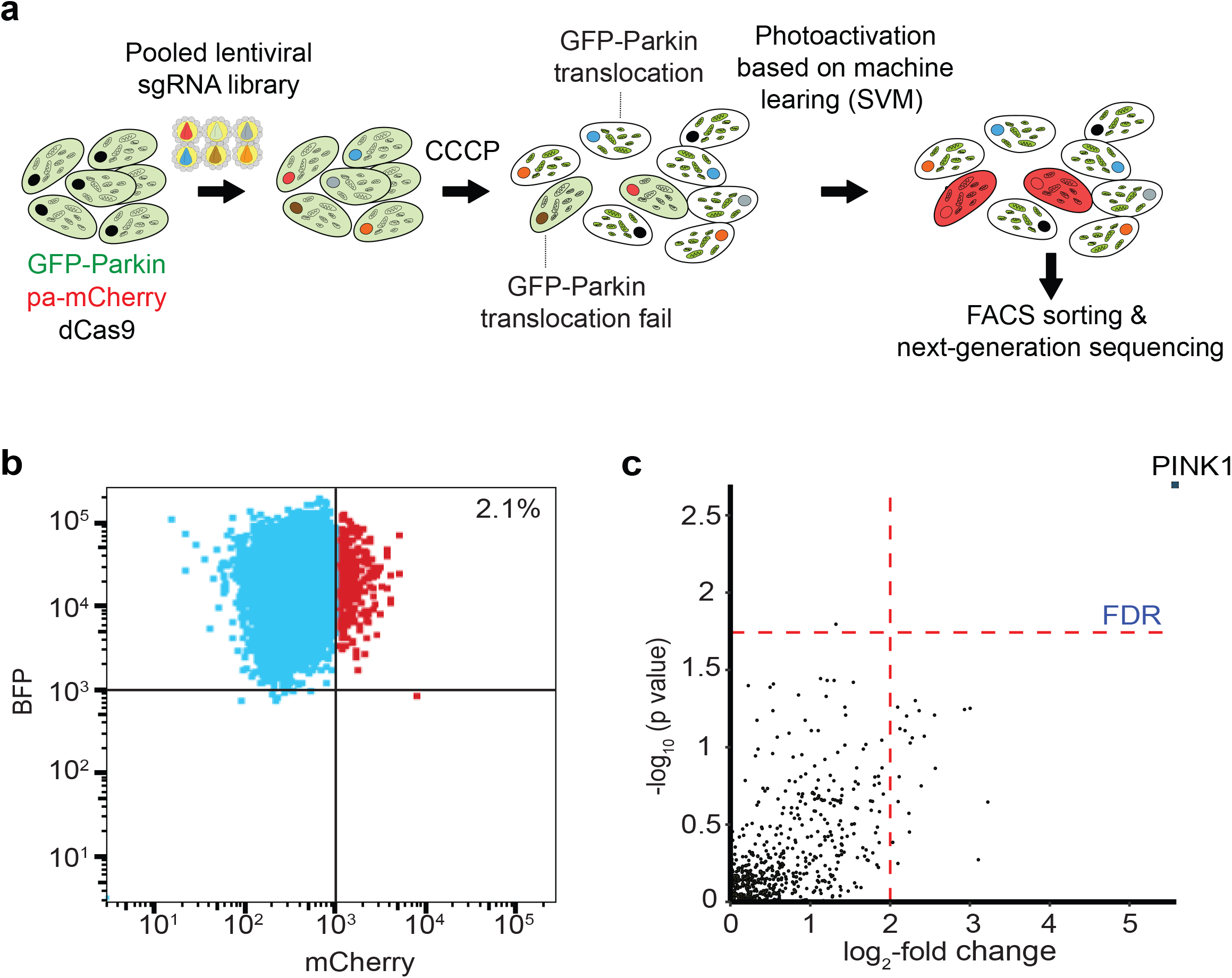
**Validation of the platform with a Parkin localization screen**, targeting Kinases, Phosphatases, and Drug Targets - gRNA pooled library (12,500 sgRNAs targeting 2774 genes). **(a)** Schematic representation of AI-PS platform **(b)** Flow cytometry scatterplot representing the separation of the post-screen photoactivated (mCherry florescence signal x-axis in red) from the inactivated cell population (BFP florescence signal y-axis in cyan). **(c)** Enrichment plot comparing sgRNA abundance in the photoactivated sample follow CCCP treatment to sgRNA abundance prior to treatment. Vertical red line set on log2-fold change threshold; horizontal red line indicating the Benjamini-Hochberg corrected p-value set on 5%. See also Table S1.

### TFEB nuclear localization screen: convolutional neural network (CNN) based screen

To explore a subcellular phenotype with more complex regulation, we screened for genes affecting the nuclear localization of the transcription factor, TFEB. Upon nutrient starvation, TFEB moves from the cytosol to the nucleus, where it activates the transcription of lysosome- and autophagy-related genes (Settembre et al., 2011). Upon prolonged starvation, mTOR is reactivated, presumably due to replenishment of nutrients through autophagy, lysosomes repopulate the cells (Yu et al., 2010) and TFEB returns to the cytosol (Fig. 4 b, Fig S4 and Video 2). As the import of TFEB to the nucleus is well elucidated (Puertollano et al., 2018), we assessed TFEB reappearance in the cytosol following prolonged starvation-induced nuclear import. U2OS cells stably expressing GFP tagged TFEB, PA-mCH, and dCas9 (designated as TFEB-GFP) were infected with a lentiviral library expressing sgRNAs against the entire genome divided in seven separate subpools (Horlbeck et al., 2016). The screen was split into seven subscreens, one per day for seven days. To increase reproducibility, each subpool screen was repeated at least 3 times.

**Figure 4.**
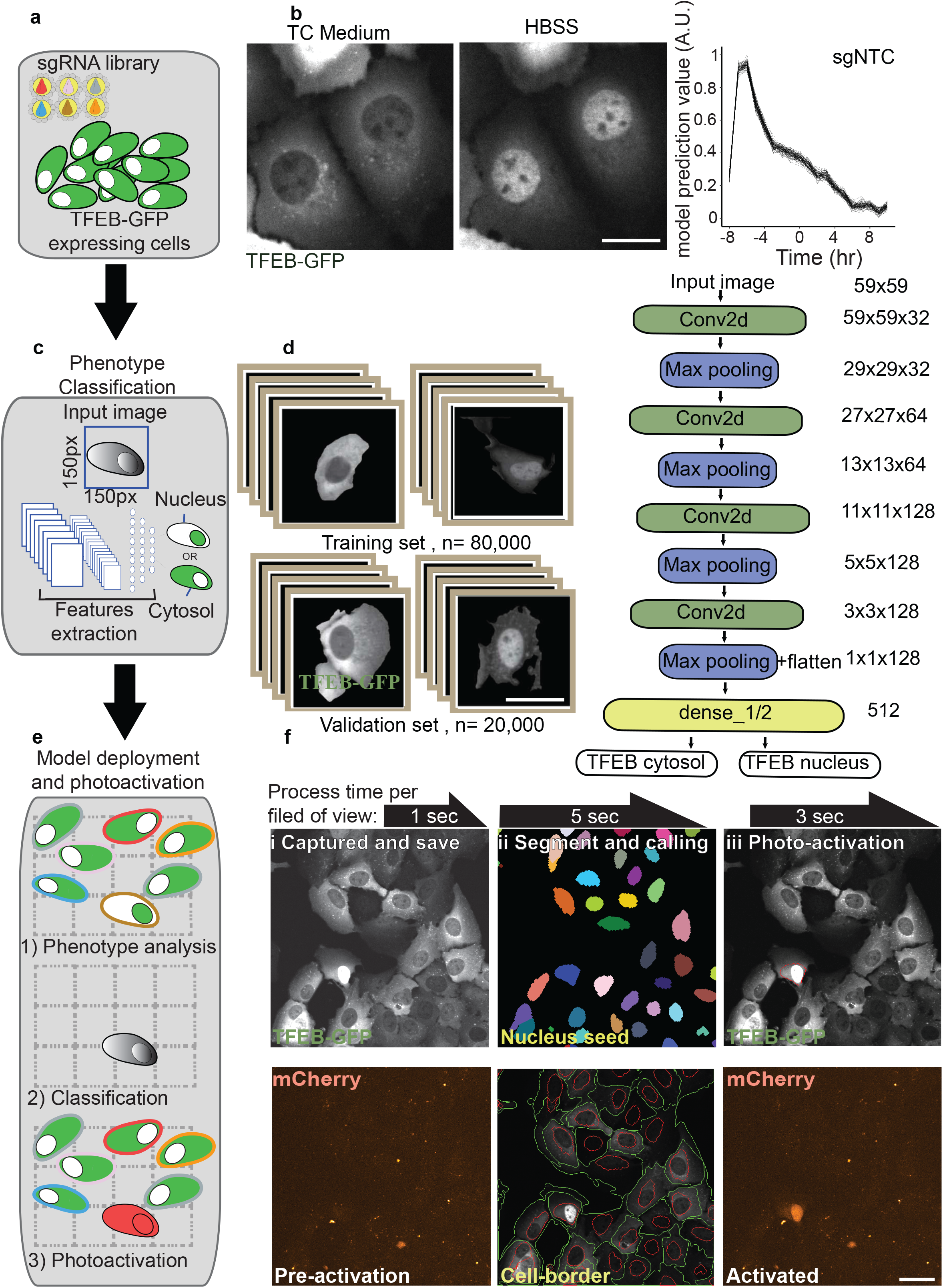
Deep learning genetic screening platform for TFEB localization. **(a)** Pooled sgRNAs infecting target cells. **(b)** Nuclear TFEB-GFP in cells treated with HBSS (1 hr) or cytosolic in cells in complete medium (cm); scale bar, 5 μm. **(c, d)** Learning set composed of 100,000 single-cell images was used for convolutional neural network (CNN) classification. ImageNet like CNN architecture is composed of four sets of convoluted processes followed by the max pooling procedure. The phenotype decision is based on probability value. Low probability value is assigned to cells with cytosolic TFEB-GFP and high probability values for cells containing nuclear TFEB-GFP. **(e)** Illustration of optical decision making coupled with photoactivation. **(f)** Representative Field Of View (FOV) of the TFEB-GFP U2OS cell screening procedure. Images were captured and saved on a local computer. i. Cell borders were identified (green circle surrounding cell border, red circle nucleus) following ii nuclear segmentation (color labeled nuclei). iii CNN classification model was deployed and masked (red circle) and photoactivation of the CNN identified cell; scale bar, 10 μm.

Because SVM classification failed to predict TFEB nuclear localization accurately (performance compression between area under the precision-recall curve of 72% for the TFEB SVM classification model versus 99% for the Parkin model (Fig. S2 c and Fig. S3 a), we used deep learning via a convolutional neural network (CNN) (Fig. 4 c,d). The training set was composed of 100,000, 150 pixel X 150 pixel single-cell images. The CNN architecture was based on ImageNet (Deng et al., 2010) architecture and composed of three deconvolutions and four Max pooling processes, which were followed by a fully connected dense network (Fig. 4 c,d).

TFEB-GFP cells expressing guide libraries were grown under complete nutrient deprivation conditions for eight hours prior to the commencement of screening after which those cells retaining TFEB in the nucleus were photoactivated (Fig. 5 a), isolated by FACS and deep sequenced (Fig. 4 e,f and Fig. S3b, Video 3). Among the seven subpooled libraries, a mean accuracy of 90% was calculated from the approximation of the area under the precision-recall curve (Fig. S3 c). The entire photoactivated and sorted gene abundance ranking list analysed for ontology clusters revealed enrichment in mitochondrial and kinase complex gene sets (Puertollano et al., 2018; Nezich et al., 2015), that may relate to energetic consequences to mitochondrial states and TFEB post-translational regulation, respectively (Fig. 5b and Fig. S5). Plasma membrane proteins were also enriched, perhaps related to cell division rates or nutrient import.

**Figure 5.**
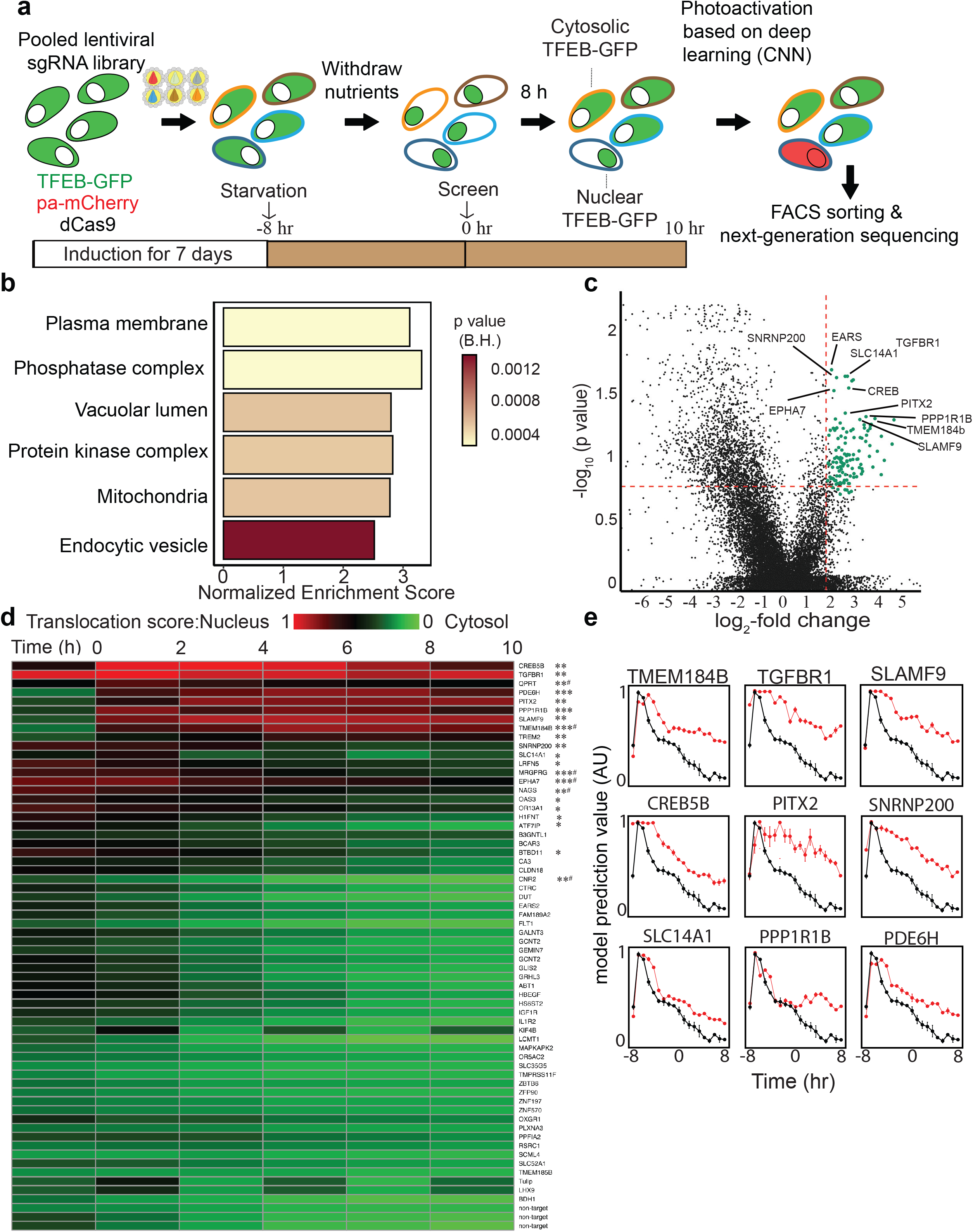
Whole-genome TFEB-GFP localization screen. **(a)** Screen workflow, TFEB-GFP cells were transduced with pooled sgRNA libraries targeting the whole genome for seven days. Following eight hours of starvation, AI-PS screening platform was initiated. **(b)** GSEA pathway analysis annotated using the gene sets derived from the GO Cellular Component database of the Molecular Signatures Database. On the x-axis is the GSEA normalised enrichment score and the color of the bars represents the GSEA calculated FDR probabilities. **(c)** Volcano sgRNA plot comparing sgRNA abundance in the photoactivated samples following eight hours of starvation to sgRNA abundance in the unsorted sample. Vertical red line is set at the log2-fold change threshold; horizontal red line is set at the Benjamini-Hochberg corrected p-value of 15%. Genes selected for secondary screening are shown in green. See also Supplementary Table 2. **(d)** Top candidates from the primary screen were selected for secondary screening. The TFEB-GFP localization CNN model probability value was used for measuring the perturbation effect on TFEB-GFP localization over time during starvation. Heat map including all genes included in secondary screen; low probability values are shown in green for cytosolic TFEB-GFP, high probability values (red) for nuclear TFEB-GFP. n=3, *p < 0.05, **p < 0.01, or ***p < 0.0001 obtained using repeated measures Anova test. p value, one sgRNA was significant. e. TFEB-GFP translocation dynamics observed during starvation for selected gene candidates, non-targeted sgRNA in black, gRNA target the designated protein in red. Quantification is displayed as mean ± s.e.m. from 3 independent experiments.

Differential sgRNA abundance analysis between unsorted and photoactivated/sorted samples showed a significant fold-change enrichment in 64 genes (Fig. 5c, Table S2). A second validation screen was conducted of the 64 enriched genes using two new sgRNAs. As with the whole-genome screen, TFEB-GFP nuclear localization following validation guide transduction during prolonged starvation was recorded 8 hours after starvation for 10 hours. The perturbation effect on TFEB positioning was compared to a non-targeting control sgRNA. We found that 21 of the 64 sgRNAs from the whole genome analysis significantly extended nuclear TFEB retention (p < 0.05, repeated-measures ANOVA, Fig. 5 d). Interestingly, these 21 validated hits were amongst the genes with the highest ranked p-value significance in the whole genome screen (Table S2). Amongst the validated genes, the signaling receptor TGFBR1 was enriched in the secondary TFEB screen (Fig. 5 d,e and Fig. 6a, Video 4). This may be related to a previous report of the induction of another MITF family transcription factor, TFE3, by the loss of TGFBR1 (Sun et al., 2016). Validating the screen, one of the strongest hits is the transcription factor, CREB (Fig. 5d,e and Fig. 6b, Video 5), which has been shown previously to mediate autophagy and induce the expression of several autophagy genes including Ulk1, Atg5 and Atg7 upstream of TFEB following starvation and TFEB itself (Seok et al., 2014). Certain autophagy genes are more predominantly activated by CREB and others more by TFEB. In addition, loss of another hit, Pitx2, *in vivo* causes an increase in mitophagy that has been linked to TFEB activation (Nezich et al., 2015; Chang et al,. 2019). Additionally, the membrane protein, TMEM184b, has been reported previously to play a role in autophagy (Fig. 5 d,e) (Bhattacharya et al., 2016; Agod et al., 2018). Loss of the phosphatase, PPP1R1B, which also scored amongst the top validated hits, resulted in significant retention in TFEB in the nucleus upon starvation (Fig. 5 d,e and Fig. 6c, Video 6). As phosphorylation of TFEB is intimately linked to its activation and subcellular localization (Puertollano et al., 2018), this hit deserves further mechanistic study. The extensively studied TFEB regulator mTOR was not significantly enriched in our photoactivated samples. To explore this explicitly, live-cell imaging of starved GFP-TFEB infected with two distinct sgRNAs targeting mTOR showed an accumulation of TFEB on lysosomes, which resulted in punctate cytosolic foci, similar to previous reports (Martina and Puertollano, 2013; Settembre et al., 2012) (Fig. 6d, Video 7). Therefore, mTOR was not identified in the enrichment analysis owing to the lack of classification of this specific mTOR phenotype, which is distinct from the deep learning model trained for nuclear localization.

**Figure 6.**
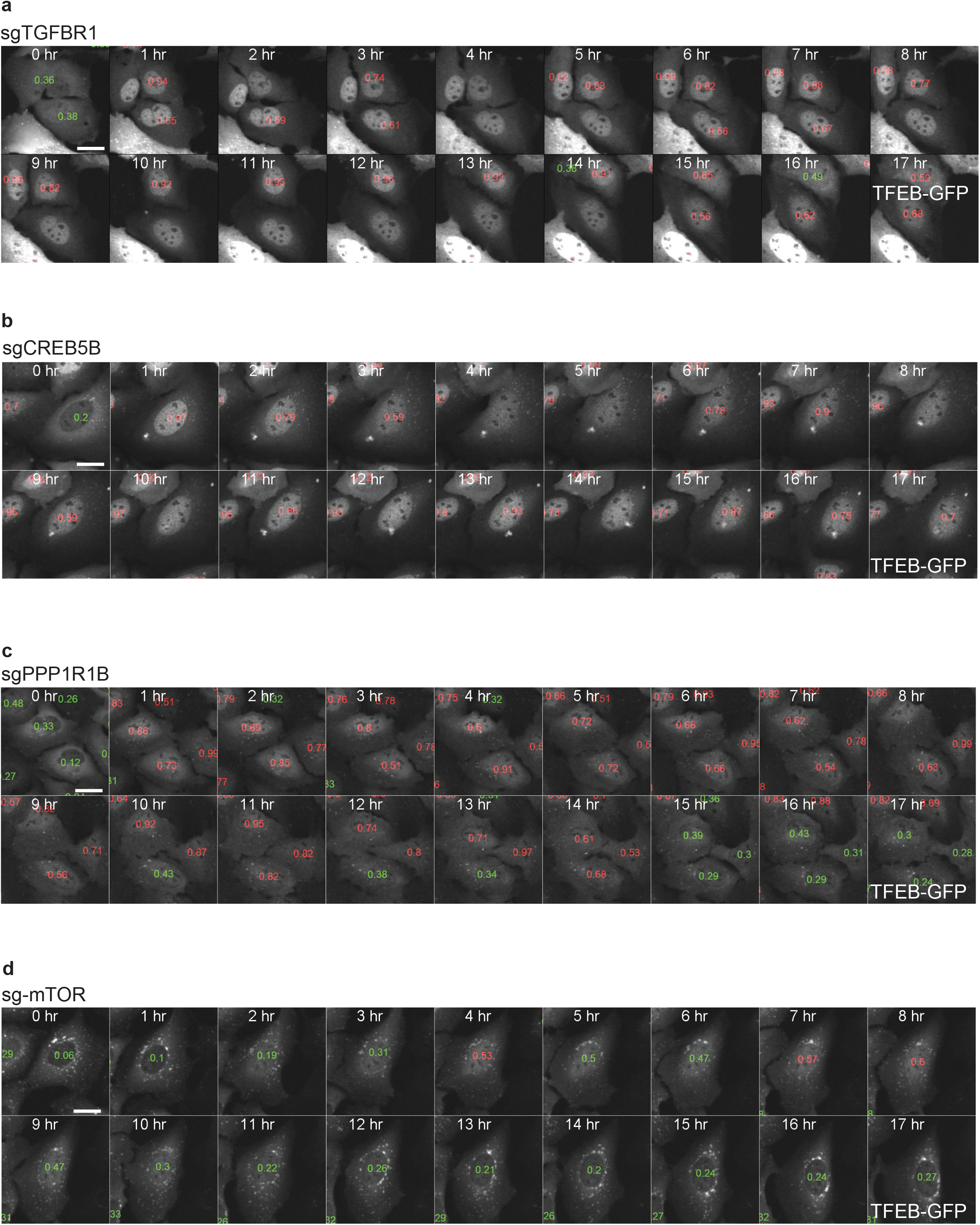
Selected sgRNA targeting genes validated in the TFEB-GFP secondary screening. low probability values are shown in green for cytosolic TFEB-GFP, high probability values (red) for nuclear TFEB-GFP. TFEB-GFP-expressing U2OS cells treated with the designated sgRNA were starved in HBSS for 18 hours, and images acquired every hour. scale bar 5 μm.

## Discussion

Here, we present a platform that applies machine learning and deep learning algorithms to allow pooled genetic screening for subcellular image phenotypes. This method, which we call Artificial Intelligence-Photoswitchable Screening (AI-PS), reduces the time, cost and complexity compared to standard screening methods that have required arrayed RNAi or CRISPR libraries.

The speed of AI-PS screening relies on the simultaneous execution of four steps: image capture, segmentation, generation of classification region of interest, and photoactivation of the region of interest. For a field of approximately 200 cells, these four steps together take an average of 10 seconds, which is then iterated across an entire plate. Therefore, a screen of 600,000 cells infected with 1/7^th^ of the genome guide library, composed of 12,500 sgRNAs, takes ~12 hours. Hence, this accelerated platform coupled with a user-friendly interface should accelerate the utility of pooled genomic screens.

The method enables the detection and labelling of cells according to subcellular protein localization. We validated this by identifying PINK1 as the only significant hit required for Parkin translocation to damaged mitochondria within the genome guide sub-library of kinases, phosphatases and the druggable genome, demonstrating an exceptional signal-to-noise ratio when using the method.

We also applied AI-PS to explore a completely different protein translocation process, one which again would be undetectable via FACS separation of whole cells based on a change in overall fluorescence intensity. The transcription factor, TFEB, is retained in the cytosol in growing cells and upon starvation relocalizes to the nucleus, where it induces transcription of lysosomal- and autophagy-related genes (Settembre et al., 2011; Sardiello et al., 2009). Upon prolonged starvation, TFEB returns to the cytosol via an undefined process. Either nuclear TFEB migrates back to the cytosol or nuclear TFEB is degraded while newly synthesized TFEB repopulates the cytosol. As we found minimal evidence for a role of cytoskeletal or nuclear transporter proteins, whether the appearance of TFEB in the cytosol is due to physically shuttling of preexisting TFEB or to an increase in translation of new TFEB remains an open question. Beyond protein localization screens, our method will be useful to identify genes involved in regulation of organelle abundance, size and shape.

Similar to the concepts presented in our study, machine learning-based image analysis has been used for calling and sorting of cells (Ota et al., 2018; Nitta et al., 2018). However, AI-PS conveys distinct advantages. First, the microscopic resolution of AI-PS is much higher than that utilized during dissociated cell sorting (Ota et al., 2018; Nitta et al., 2018), allowing the identification of more difficult to detect subcellular structures. Specifically, the detection of minor subcellular events such as alteration in protein distribution, positioning and motion requires high spatial-temporal resolution image acquisition. Previously published methods utilized low magnification objectives (4x and 10x) and very short exposure times (below 50 msec), which resulted in low signal-to-noise ratios, and are not suitable for the resolution of subcellular events.

Another advantage of the AI-PS platform is its wide accessibility – there is no need for specialized flow instrumentation, and the algorithms and code presented here can be adapted easily for a variety of microscope systems. AI-PS is compatible with adherent tissue culture cells, unlike sorting-based approaches for which cells must be in suspension, further allowing a more accurate examination of subcellular events in regular culture conditions. One current limitation of AI-PS is that the cells must be screened live to allow trypsinization to produce single cell suspension for FACS. Because some phenotypes would be better screened in fixed cells, we are developing methods enabling single cell release of fixed cells to allow screening of additional cell biology processes.

Another step to improve AI-PS would be to reduce the segmentation time per image to speed up the screen. Fortunately, a huge improvement in cell segmentation, specifically the development of deep learning-based techniques, such as U-Net segmentation (Caicedo et al., 2019; Toth, 2018; Ronneberger et al., 2015) can be utilized in AI-PS. This new deep learning-based segmentation has the potential for at least a five-fold reduction in analysis time. Increasing the speed will make it possible to increase the sample size and thereby increase the sgRNA coverage in the sorted samples, decreasing the false discovery rate. Another strategy for increasing specificity would be to utilize single-cell DNA sequence analysis.

In conclusion, our platform demonstrates novel implementation of machine learning to improve cell biology research and discovery and enables phenotypic-based screening at the subcellular level – an approach previously largely unavailable. Additionally, AI-PS can be implemented for drug target exploration and may prove valuable in methods targeting single cells within complex human samples.

## Methods

### Cell lines, constructs and reagents

U2OS and HEK293T were cultured in a humidified incubator at 37°C and 5% CO_2_ and maintained in DMEM (Life Technologies) supplemented with 10% (v/v) FBS (Gemini Bio Products), 10 mM HEPES (Life Technologies), 1 mM sodium pyruvate (Life Technologies), 1 mM non-essential amino acids (Life Technologies) and 2 mM glutamine (Life Technologies). Testing for mycoplasma contamination was performed bimonthly using the PlasmoTest kit (InvivoGen).

For constituting a stably expressing dCas9-KRAB U2OS cell line we took a similar approach as was described here. In brief, pC13N-dCas9-BFP-KRAB (127968, Addgene) was integrated in the U2OS genome using F-Talen and R-Talen (pZT-C13-R1 and pZT-C13-L1, Addgene:62196, 62197) targeting the human CLYBL intragenic safe harbor locus between exons 2 and 3 (as was described previously (Tian et al., 2019)). The U2OS-dCas9 cell line was then subcloned and the dCas9 activity assessed to select the most potent clones for further use (Fig. S1 c).

To generate the parental U2OS-dCas9-PA-mCh, photoactivatable-mCherry was PCR-amplified from the plasmid N-PA-mCh and assembled into the retroviral vector pBABE-puro using HiFi DNA Assembly (E5520S, NEB). To create the stable U2OS-dCas9-PA-mCh/GFP-Parkin and U2OS-dCas9-PA-mCh/TFEB-GFP cell lines, Parkin or TFEB was inserted into the lentiviral pHAGE vector by HiFi DNA Assembly (E5520S, NEB). The cell lines were subcloned and cells expressing low levels of the GFP-tagged proteins were selected to prevent overexpression artifacts. For nucleus segmentation we used a lentiviral plasmid expressing nuclear-localized Halo-tag, hU6-bsd-NLS-Halo. Prior to the screen, HBSS was supplemented with 2 μM of the pa Janelia Dye 646, SE, (TOCRIS). For the Parkin screen, the nucleus was detected using 1000x dilution of Draq5 (62251, ThermoFisher).

For Parkin-induced mitophagy, GFP-Parkin cells were treated with 10 μM Carbonyl Cyanide Chlorophenylhydrazone (CCCP) (Sigma-Aldrich) and 0.1 μM Bafilomycin A (Sigma-Aldrich). For TFEB screening, cells were starved in Hank’s Balanced Salt solution (HBSS) without calcium and magnesium (14170112, ThermoFisher).

### Parkin-GFP and TFEB-GFP positioning classification by support vector machine (SVM)

To create the classification model, we initially trained 2,234 images of each of the binary phenotypes: Parkin or TFEB translocation. GFP-Parkin signal was mitochondrial versus cytosolic while TFEB-GFP was nuclear versus cytosolic. The model was created using the R library e1071. In brief, we used a radial basis Kernel with a cost violation of 10 computed for an example set of phenotypes using the radial Kernel formula: e^(−γ|u−v|^2)^ For optimization of the model, we performed iterations and calculated performance by area under the receiver operating characteristic (ROC) curve or precision-recall curve (in the case of asymmetric phenotype representation). The performance values were plotted against iteration to prevent data overfitting.

### GFP-TFEB positioning classification by Convolutional neural network

For TFEB localization classification, an ImageNet (Deng et al., 2010) architecture CNN model was created using TensorFlow and the KERAS library. A training set composed of 107,226 single-cell example images of GFP-TFEB in the nucleus or cytosol was produced. 80% of the data was used for training and 15% for validation. The remaining 5% was used for testing the model performance. Image input size was 150 pixel X 150 pixel and three steps of convolution and max pooling were conducted at a learning rate of 1e^−4^.

Training was performed with 50 epochs and a batch size of 200. Overfitting was prevented by employing the built-in Keras callbacks API feature to save the model weights after each epoch. The selected model was chosen from the epoch* at which the validation and training loss curves were no longer decreasing. The variation in fluorescence signal intensity was accounted for by randomly applying brightness augmentation (10% to 90%) to the images in the training data set.

### Model performance

To assess classification model performance, we performed a precision-recall curve in which the curve integral was a measurement of accuracy (Fu and Yi, 2019). In brief, 5-10% of images in the data set from our experiment were arbitrary selected for performance testing. Images were collapsed into single cells. The parameters extracted for constructing the precision-recall curve were the corresponding CNN prediction value against the ground truth class. The curve and the AUC were plotted and calculated using the R package PRROC (Grau et al.).

### Image acquisition and model deployment

SVM deployment live-image acquisition was done on a Nikon Ti-2 CSU-W1 spinning disk confocal system equipped with a high-speed electron-multiplying charge-coupled device camera (Evolve 512; Photometrics) using a 20X air objective (NA 0.75) with an environmental control chamber (temp controlled at 370C and CO2 at 5%) operated by Nikon elements AR microscope imaging software.

Cells were seeded at for screening at 1×10^5^ cells per well of a 2 well Lab-Tek chamber slide (Thermofisher, 155360). The on-the-fly real-time capture was done using the 488 nm laser channel for excitation and using the 520 nm emission detector to collect the GFP signal and the 647 nm excitation laser and 667 nm emission detector for the segmentation channel. Saved images were segmented live using a bash file script (https://github.com/gkanfer/AI-PS), and the classifications were deployed by the SVM model. A mask file containing the selected cells was generated and stored on the local computer. The mask image was used to photoactivate the called regions by exciting with a 405 nm wavelength using a Bruker minscanner XY galvo photostimulation scanner. The process was iterated across more than 1000 fields of view (512 px by 512 px for the Parkin screen and 2048 px by 2044 px for the TFEB screen). The Nikon Imaging System (NIS) elements AR microscope software was used in JOB mode to allow integration of the deployment code on-the-fly (the JOB file can be found on our https://github.com/gkanfer/AI-PS/). In brief, following capture and saving of the 488 nm and 647 nm channel images on the local computer, the NIS JOB module OUTPROC is activated and directed to run the segmentation and deployment R script. Next, the region of interest mask is generated, uploaded back on the local microscope computer hard drive on the outproc folder path, after which the NIS-JOB continues by saving the mask coordinates and preforming the photoactivation of the selected regions of interest with a 405 nm laser. The microscope stage then moves to the next field of view to repeat the process.

Live-cell image acquisition and deployment of the CNN-based screen were carried out on the Eclipse Ti2-E (Nikon) with CSU-W1 spinning disk system equipped with an ORCA-FLASH 4.0 V3 sCMOS (Hamamatsu) and an Opti-Microscan XY Galvo Scanning Unit and a Nikon LUN-F laser unit with 90mW 405NM, rated 90mW output at fiber tip, using a 20X objective (NA 0.75) and environmental control chamber (temp controlled at 37°C and CO_2_ at 5%). The microscope was controlled by the NIS elements AR microscope imaging software.

The on-the-fly real-time acquisition and deployment for the CNN-based screen were performed as described above with one major modification: the TensorFlow deployment script was running the backend “while-loop” throughout the acquisition (https://github.com/gkanfer/AI-PS/tree/master/TFEB_screen).

### Cell segmentation analysis and processing

For image manipulation, the R package EBimage (Pau, 2009) was used similar to a previous report (Laufer et al., 2013). In brief, the two-channel images were min/max-normalized and nuclear staining was used as a seed to identify individual cells. For nucleus segmentation, thresholding with a 5×5 filter map and Watershed transformation were applied. Then, the target channel (designated GFP), was used to identify cell borders and edges for segmentation, after which it was used for classification. High-pass filtering and local thresholding followed by global thresholding were used to create global and local masks. Together with the nucleus mask generated in the first step, this mask was used for the Cellprofiler (Carpenter et al., 2006)-based EBimage propagation function. To handle outlier cells, several features were computed and the outlier features were removed. For SVM classification, preselected features were computed and used for classification. For the CNN classification, single cells were extracted and stacked into tensor array configuration which is compatible with CNN-based prediction analysis.

### sgRNA lentiviral production

To generate lentivirus expressing sgRNA libraries, CRISPRi sub-pooled libraries were used (Horlbeck et al., 2016). On day 0, 7.5 × 10^7^ Hek293-lentiX cells (Clontech) were seeded on 15-cm tissue culture plates. The next day (Day 1), 20 μg/ml subpooled sgRNA plasmid, 14.1 μg/ml PAX2, 4.2 mg/ml MDG2 and 1.2 μg/ml pAdvantage (3rd generation lentiviral vector packaging systems) were transfected using 75 μl of Lipofectamine 2000 (11668019, ThermoFisher) in Opti-MEM (ThermoFisher). On day 2 media was changed, and on day 3 virus was harvested. A lentivirus precipitation kit (VC100, Alstem Cell Advancements) was used according to the manufacturer’s suggestions to concentrate the virus.

To determine MOI, 0.1 × 10^6^ cells were seeded in 24-well plates and infected with four titrations of the concentrated virus. Genomic DNA was isolated using QIAamp DNA Micro Kit (56304, Qiagen). The number of genomic viral integration sites was compared with the number of housekeeping genes using a ddPCR—BioRad QX200 AutoDG Droplet Digital PCR System (BioRad). The volume to MOI ratio was calculated using the following formula:

1. Insertion number (from ddPCR) × dilution factor = Transducing Units (TU)
2. (Desired MOI × cell number) / Transducing Units = Virus Volume

The ddPCR primer mix for amplifying upstream of the sgRNA integration region was purchased from BioRad: Gaagaagaaggtggagagagagacagagacagatccattcgattagtgaacggatcggcactgcgtgcgccaattctgcagacaaatggcagtattcatccacaattttaaaagaaaagggggg (FAM) The house keeping probe used for comparison was EiF2C1 (Assay ID: dHsaCP2500349 Cat: 10031243, BioRad).

To conduct the screen, library expression, 5 × 10^6^ dCas9-PA-mCh-expressing cells were seeded on day 0. The next day, the appropriate virus volume was added to cells to achieve an MOI below five. Two days after infection, sgRNA-expressing cells were sorted using a 407 nm Laser and 450/50 nm filters. Following four days of growth, cells were reseeded in 2-well screening chambers. To maintain sufficient sgRNA representation, cells were maintained at numbers corresponding to a coverage of at least 100 cells per sgRNA.

### Activated sample isolation

After screening, cells were detached using trypsin (Sigma), washed once with PBS and filtered using a 50 micron sieve (Corning) to obtain a single-cell suspension. The volume was adjusted to obtain up to 10 million cells per ml using PBS. Cells were kept in the dark on ice until sorting, which was done using a BD FACS Aria cell sorter equipped with 355 nm, 407 nm, 532 nm, and 640 nm laser lines and BD FACSDIVA software to perform aseptic cell sorting. Physical properties (FSC and SSC parameters) of cells were used to identify and exclude debris, dead cells and doublets. All single cells were then selected for GFP expression using the signal from the 488 nm laser line 515/30 nm filters. mCherry signal was identified using the 532 nm laser line and 610/25 nm filter and BFP signal was identified using signal from the 407 nm Laser and 450/50 nm filters. Cells were purified into two populations, which were either GFP+/BFP+/RFP+ or GFP+/BFP+/RFP-for downstream analysis.

### Illumina library construction and sequencing

Following FACS sorting, samples were pelleted by centrifugation and subjected to genomic DNA isolation using the QIAamp DNA Micro Kit (56304, Qiagen). To construct the sequencing library, genomic DNA was amplified by two-step PCR. In the first step, Unique Modifier Identifiers (UMI) fused with lentiviral vector integration site (step 1 Fw primer) were mixed with 7i adaptor primer fused with lentiviral vector integration 3’prime integration site (step 1 Rev primer). The mixture was amplified using 5-10 PCR cycles. The second amplification step included a forward primer complementary to the UMI primer fused to 5i (step 2 Fw primer) Illumina adaptor primers and 7i (step 2 Rev primer) and amplified using 25 PCR cycles. DNA concentration were measured using the NEBNext® Library Quant Kit for Illumina (E7630L, NEB). Each 50 μl PCR reaction was composed of 0.5 μM primers, 0.5 μl of Phusion hot-start DNA polymerase (F549S, ThermoFisher) and 2.5 μM dNTPs (N0447S, NEB). After 25 cycles (second PCR step), the PCR products were cleaned using AMPure beads (A63880, Beckman Coulter) according to the manufacturer’s protocol.

Fragment size and purity was determined using Agilent TapeStation 2200 and 4200 models, and the desired fragment size of 300 bp was extracted and eluted using a Pippin instrument (Sage Science) with HT 2% Agarose Gel, 100-600 bp (HTC2010). For the Parkin screen, we used 300 v2 Cassettes (15 million reads) on MiSeq (MS-102-2002), whereas, for the TFEB screen, Illumina pair-end sequencing was performed on a NextSeq 550 instrument using a sequencing chip of 300 Mid Output Kit v2.5 (120 million reads, cat 20024905, Illumina). The read length was 200 bp and 7 bp for the indexing primers. Custom sequencing primer were used: (UMI sequence-N, Index sequence-n).

Primer set for step 1:

Fw: 5’-AAGCAGTGGTATCAACGCAGAGTAC**NNNTNNNTNNNTNNNNNNNN**GCACAAAAGG AAACTCACCCT-3’
Rev: 5’-CAAGCAGAAGACGGCATACGAGAT**nnnnnnn**CGACTCGGTGCCACTTTTTC-3’

Primer set for step 2:

Fw: 5’-AATGATACGGCGACCACCGAGATCTACACAAGCAGTGGTATCAACGCAGAGTAC-3’
Rev: 5’-CAAGCAGAAGACGGCATACGAGAT**nnnnnnn**-3’
Sequencing primer: 5’-TTATCAACTTGAAAAAGTGGCACCGAGTCG-3’

### UMI extraction and read count generation

The sgRNA abundance analysis was split into four parts: first, the fastq file was demultiplexed according to the run sample sheet using the FASTX Barcode Splitter. Second, using UMI-tools the sequences were extracted and low-quality sequences were trimmed using trimmomatic. Sequences were aligned and mapped to the library data set using Bowtie and Tryhard modules as described previously (Horlbeck et al., 2016). Finally, deduplication grouping and counting were conducting using UMI-tools. The complete Unix based bash file is available on Github.

### Differential sgRNA abundance analysis

The differential abundance of sgRNAs between photoactivated-sorted samples and control untreated samples were assessed using the EdgeR package. First, samples were log2- and count-per-million normalized. Sample variation was determined by covariance-based PCA analysis and read count flooring was established by modeling the noise using coverage as a function of read count. sgRNA enrichment is defined as two standard deviations from the mean of the distribution of non-target-sgRNA controls. For gene aggregation analysis, similarly to a previous paper (Tian et al., 2019), the highest enrichment sgRNA sets were selected by bootstrapping the entire data set. Using EdgeR (Robinson et al., 2010; Dai et al., 2019), the FDR-corrected p-value was calculating by the roast function (Rotation Gene Set Test (Robinson et al., 2010)) following the exactTest function of EdgeR (n = 3 or 4 replicates). Gene set analysis was performed using GSEA 4.0.3 and our whole genome list was ranked according to FC and p-value. The pathway annotation used are the MSigDB Collections (C2:C5) (Reimand et al., 2019).

### Experimental approach for validation

For the secondary validation, the best two sgRNA with FC higher than two standard deviations from the non-targeting-sgRNA controls and roast test FDR < 15%. 128 sgRNAs targeting 64 high-scoring hits (See also Supplementary Table 2) identified from the primary pooled screen (two sgRNAs per gene) and 2 nontargeting control sgRNAs were individually cloned into the lentiviral mU6-BstXI-BlpI-BFP sgRNA vector (Horlbeck et al., 2016) and confirmed via sequencing.

Nontargeting sgRNA sequences:

Non-targeting control sgRNA 1 - 5’ GCTGCATGGGGCGCGAATCA 3’
Non-targeting control sgRNA 2 - 5’ GTGCACCCGGCTAGGACCGG 3’

To generate virus, 2 × 10^6^ Lenti-X 293T cells (Clontech) were seeded in 6-well plates in 1.5 ml DMEM (Life Technologies) supplemented with 10% (v/v) FBS (Gemini Bio Products), 10 mM HEPES (Life Technologies), 1 mM sodium pyruvate (Life Technologies), 1 mM non-essential amino acids (Life Technologies) and 2 mM glutamine (Life Technologies). Cells were transfected the next day in the following manner using Lipofectamine 3000 (Thermofisher): 1.2 μg lentiviral sgRNA plasmid, 0.8 μg psPAX2 packaging vector, 0.3 μg pMD2G packaging vector, 0.8 μg pAdvantage packaging vector, and 5 μl P3000 Reagent were dilute in 150 μl Opti-MEM and incubated 5 minutes at RT; 3.75 μL Lipofectamine 3000 Transfection Reagent (Thermofisher) was diluted into 150 μl Opti-MEM and incubated at room temperature for 5 min, after which the diluted DNA was added, mixed via pipetting, incubated at RT for 40 minutes, then added dropwise to cells. Media was replaced the next day and harvested after two days and centrifuged at 4°C for 10 min at 10,000 x g to pellet cell debris. The supernatant was aliquoted and frozen at −80°C to ensure consistency throughout the validation process.

U2OS cells expressing dCas9 and PA-mCherry were seeded at 20,000 cells per well in 96-well plates on day 0, excluding all exterior wells. On day 1, cells were transduced with virus for 24 hours with 8 μg/ml polybrene at 2 concentrations with 3 replicates per concentration, allowing 10 different viruses, including a control nontargeting sgRNA, to be tested per plate. Cells were checked visually on days 2 and 3 for confluency and blue nuclear signal indicating expression of the sgRNA. If crowded, cells were trypsinized and split to 1-2 96-well plates. Cells were split again on days 4 or 5 as needed into a 96-well imaging plate (Perkin Elmer). A half media change was performed every other day if cells were not being split. On day 7, media was removed, cells were washed 3 times and then left in warm PBS without calcium and magnesium (Thermofisher). Cells were imaged every 60 minutes for 20 hours using a 20X air objective (NA 0.75) on a Nikon Ti-2 CSU-W1 spinning disk system with a photometrics 95B camera operated by Nikon Elements software equipped with temperature regulation and CO_2_ control. For every sgRNA, 9 images per well in 3 replicate were acquired. For TFEB translocation response compression, a fixed number of single cell images (n=360) per guide RNA per time per biological repeat were normalized to nontargeted control mean value. To determine if there is a significant difference between the difference values generated for the control replicates on the same plate and the difference values for a guide’s replicates on the same plate, we used repeated-measures ANOVA.

### Sample size power calculation

To estimate the necessary screening sample size, we conducted power calculations using t-tests of means (pwr.t.test function in R). First the pooled group sample standard deviation was calculated for each sgRNA (Cohen’s d): d=sqrt((n1−1)*s1^2+(n2−1)*s2^2)/(n1+n2−2)) where n1 designated the observed number of controls and n2 the observed number of photoactivated and sorted samples. Then, d was passed into the pwr.t.test function with a p-value of 0.05 and increasing power set from 0.2 to 0.8. The pwr.t.test function yields the sample size require for detect a significant deferential abundance.

### Shiny AI-PS Application

We created a graphic user interface (GUI) in Shiny (by Rstudio) that performs each step (image segmentation and classification, and creation and testing of model) required to build and test an SVM-based classification model for AI-Photoswitchable Screening. This application can be accessed directly through the website: https://hab-gk-app.shinyapps.io/gk_shiny_app/. Alternatively, the app can be run locally from the source code found at https://github.com/hbaldwin07/GK_shiny_app. Performance is better on local machines than on the network server, so this is the recommended method for those using particularly large datasets or data files (> 10 MB per image). All instructions for running/using the program can be found on the GitHub website.

## Author contributions

G.K. led the project and performed most of the experimental work. G.K. conducted and performed all the screens and created the cell lines. K.M. and G.K. created the SVM based code. H.B. and G.K. created the Shiny APP. G.K. created the deployments and CNN code. M.K. and M.W. build the sgRNA libraries. M.K., K.J., M.W. and G.K. conducted NGS analysis and statistics. S.S. and G.K. planed and preformed the secondary analysis validation screen. G.K., S.S. and R.Y. wrote the paper. G.K., R.Y. and J.L.S. designed the study. All authors discussed the results and commented on the manuscript. R.Y. and J.L.S. supervised the project.

## Acknowledgments

We thank Nico Tjandra for intellectual contributions. We thank Nick Ader, Eric Bunker, Elyssa Hawk and Sue Smith for helping with cloning, cell lines and Lentivirus production. We thank Catherine Nezich, Hetal Shah, Jose Norbert Vargas and Benoit Kornmann for comments on the manuscript and all the Youle lab and the Lippincott-Schwartz lab members for critical comments. We thank Talya Chooly for supporting the project. Flow cytometry cell sorting and sample isolation was performed at the Flow Cytometry Core, NHLBI. Next generation deep sequencing was performed at the CCR Genomics Core, NCI. We thank the NIH based Nikon team for helping with imaging and integration of our external codes. This work used the computational resources of the NIH HPC Biowulf cluster (https://hpc.nih.gov). This work was supported by the NINDS intramural program. M.K. was supported by NIH/NIGMS grant DP2 GM119139.

---

**Figure S1.**
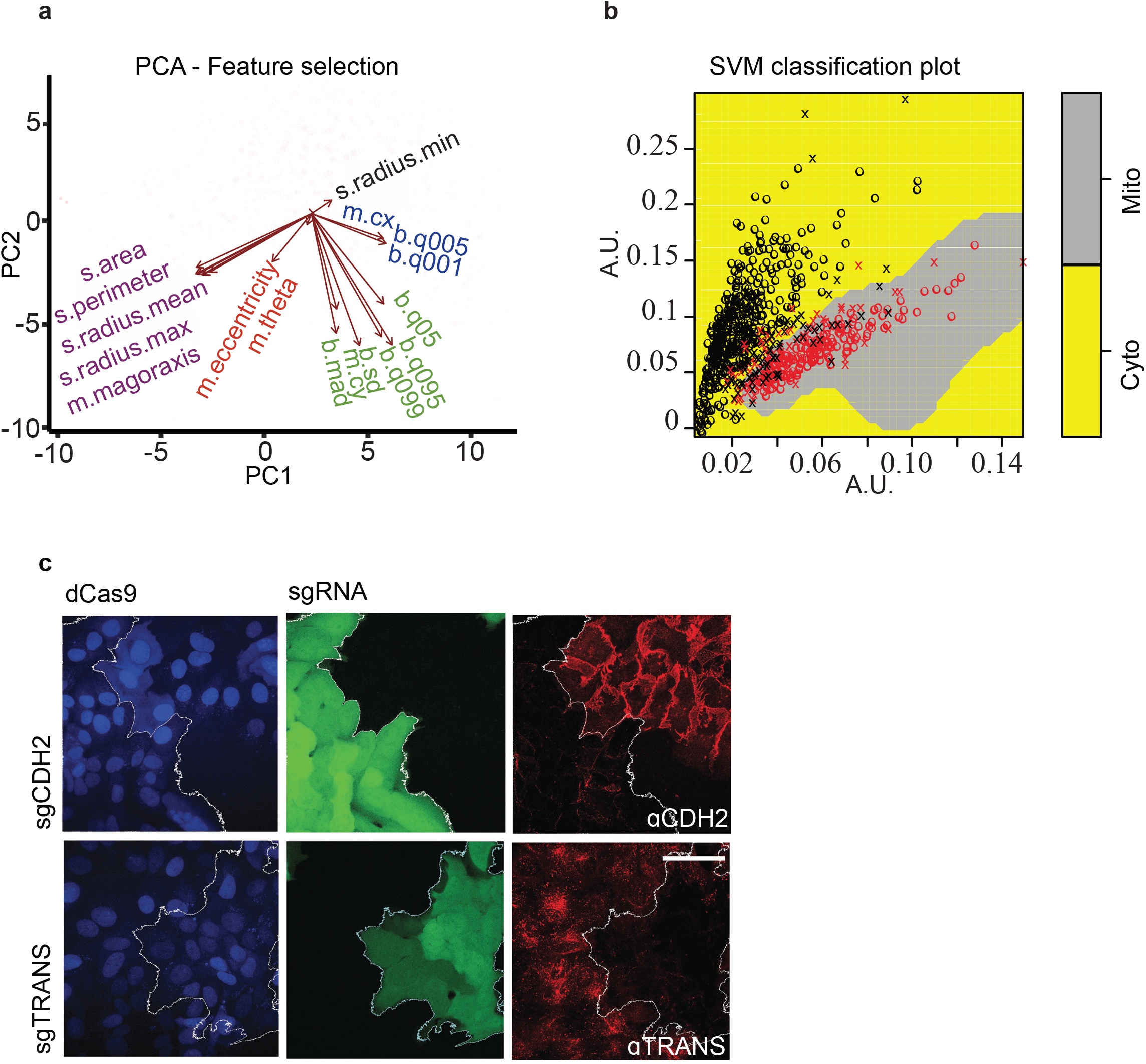
Support vector machine classification build for Parkin screen. **(a)** PCA analysis of 18 feature-predictors calculated using the R function computeFeatures from the EBImage library. **(b)** 2D representation of non-linear hyperplane separation of mitochondrial Parkin phenotype vs. cytosolic phenotype. Mitochondrial Parkin predicted cells in red, Cytosolic Parkin in black. The variable o for correct classification, and x for misclassification. **(c)** dCas9-expressing U2OS cells treated with sgRNA targeting either TRANS or CDH2 and immunostained using TRANS or CDH2 antibodies, scale bar 20 μm.

**Figure S2.**
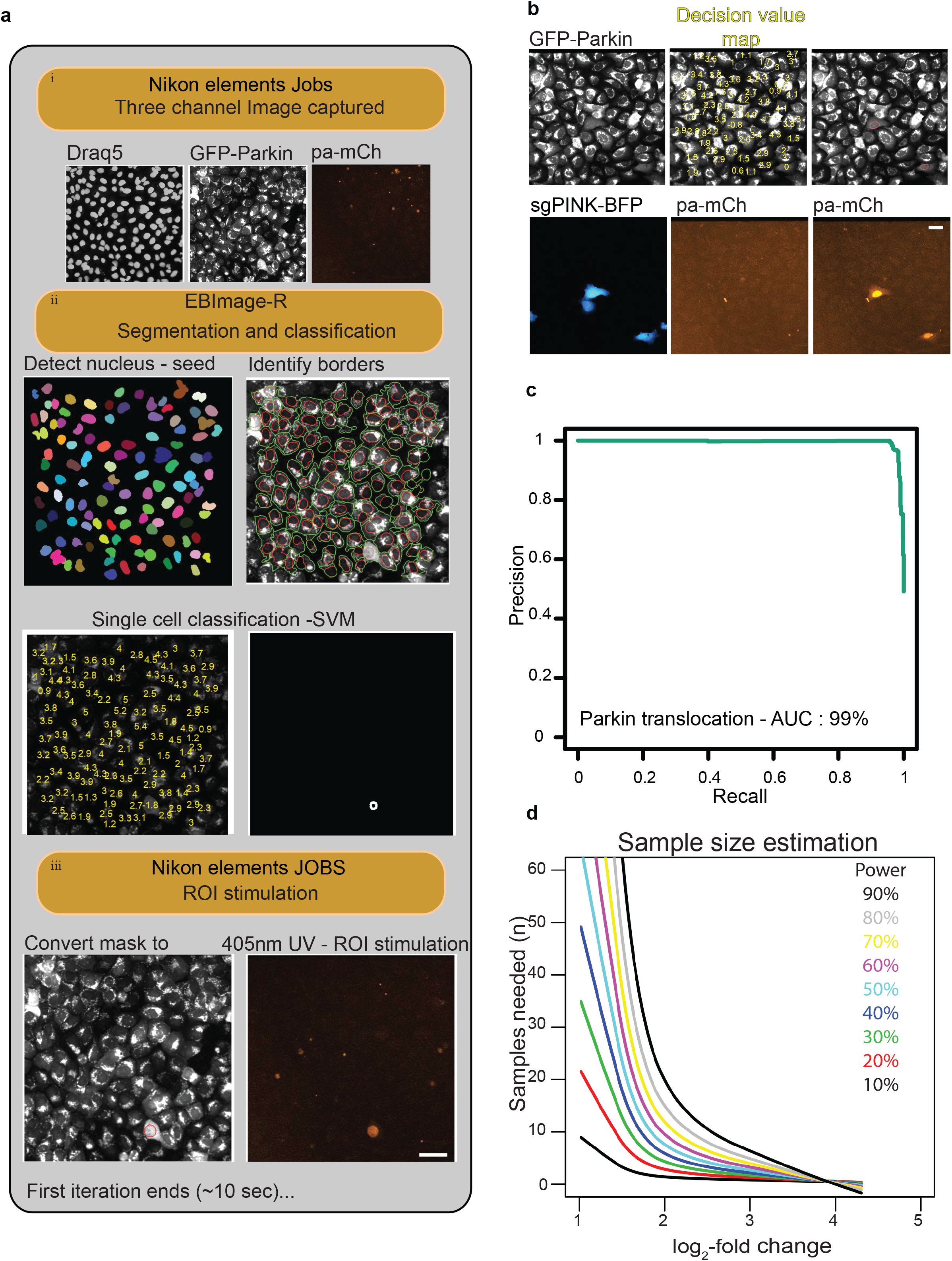
AI-PS processing stages and model assessments. **(a)** AI-PS procedure iteration on-the-fly. i. Three-channel images are acquired and available for Image processing and analysis. ii. The Draq5 channel is used for nuclear detection and the GFP-Parkin image is used to identify cell borders (right panel, red circle masks nucleus and the green circle masks the cell borders). Post-training single-cell phenotypes are scored by the SVM model (bottom left panel, decision value in yellow), where the mask is marking the predicted cells (bottom right panel, white circle). iii. Mask images are uploaded back to Nikon Elements software and converted to ROI (left panel, red circle). The ROI is stimulated by 405 nm laser for 50 ms with 100% laser power (right panel, 568 nm detector). **(b)** For proof-of-principle screening, the cell population expressing GFP-Parkin contains 10% of cells infected with sgRNA targeting Pink1 (sgRNA in blue for BFP) and treated with 1 μM CCCP; upper panel, representative GFP-Parkin images; middle panel, single-cell phenotype prediction score (decision value in yellow), score < 0.8 for cytosolic GFP-Parkin, score > 0.8 for mitochondrial GFP-Parkin; right panel, SVM-called positive cells shown on 568 nm emission detector. **(c)** Precision-Recall curve from 9,546 single cell images obtained during 12 hours live-image acquisition with 1-min intervals with 1 μM CCCP treatment supplemented with 100 nM Bafilomycin A. The accuracy is computed from the integral area under the Precision-Recall curve (AUC, Area Under the Curve). **(d)** The sample size was estimated for observing a significant effect for a log2-fold change of 1.5. The log2-fold change was modeled based on the non-targeting negative control distribution. The sample size was estimated from the GFP-Parkin screen. The effect size is estimated by the log2-fold change of the photoactivated sample vs. the control sample. The estimation is made using data gathered from 4 biological repeats of the GFP-Parkin screen. For more details see Methods.

**Figure S3.**
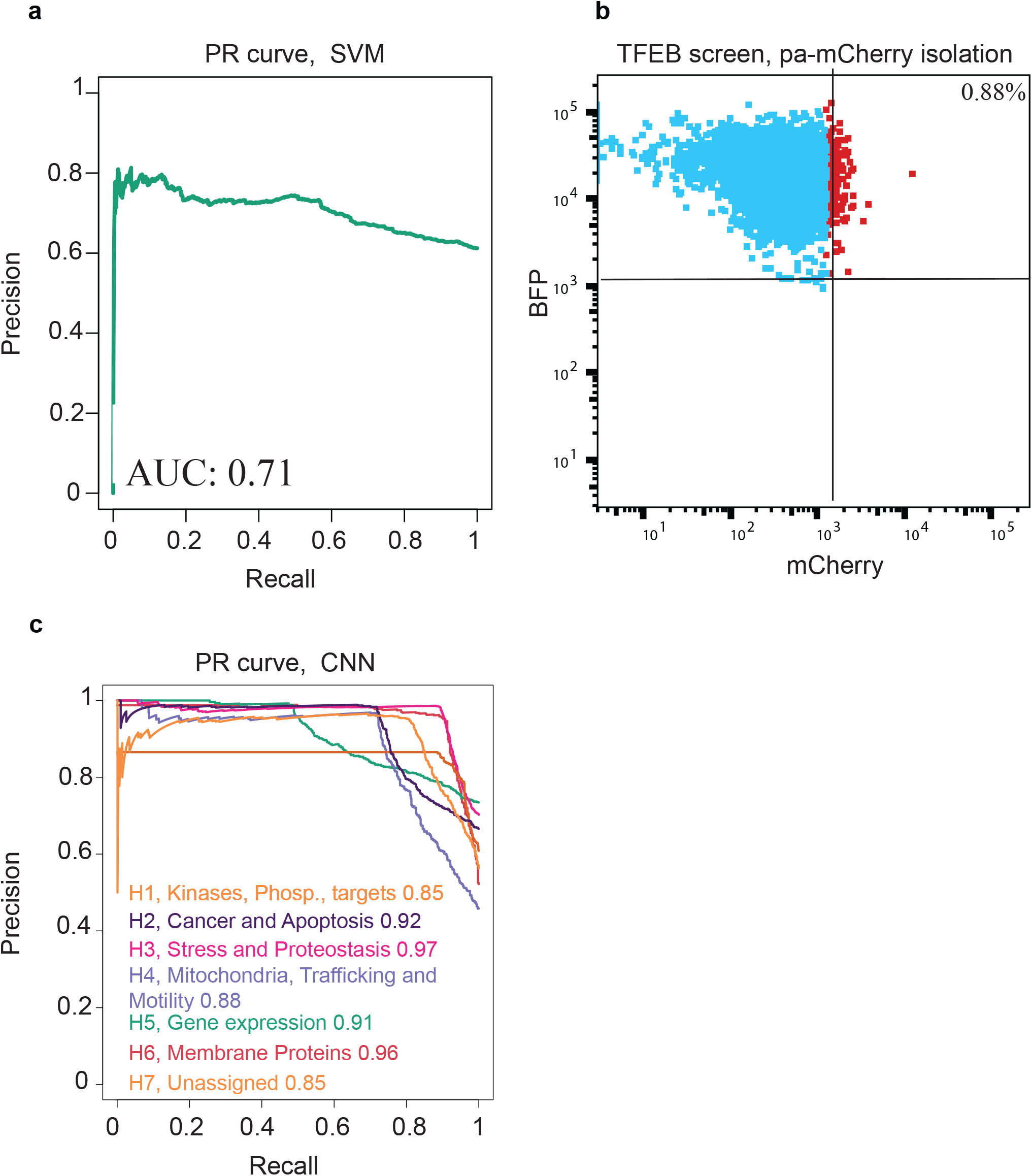
TFEB-GFP screen performance. **(a)** TFEB-GFP phenotype classification performance by SVM, Precision-Recall curve from 7,848 single cell images obtained from HBSS starved cells. Images collection began 8 hours after starvation initiated and continued for another 10 hours. Accuracy is computed from the integral area under the Precision-Recall Curve (AUC, Area Under the Curve). **(b)** Flow cytometry scatterplot representing the separation of the post-screening photoactivated mCherry signal (x-axis in red) from the inactivated cell population (BFP fluorescence signal, y-axis in cyan). **(c)** TFEB-GFP phenotype classification performance by SVM. Precision-Recall Curve from ~5,000 single cell images obtained from starved cells. Image collection began 8 hours after starvation initiated and continued for another 10 hours. The accuracy was computed from the integral area under the Precision-Recall Curve (AUC, area under the curve). The AUC was calculated per subpooled library (designated by color), from a pool of 3 biological repeats.

**Figure S4.**
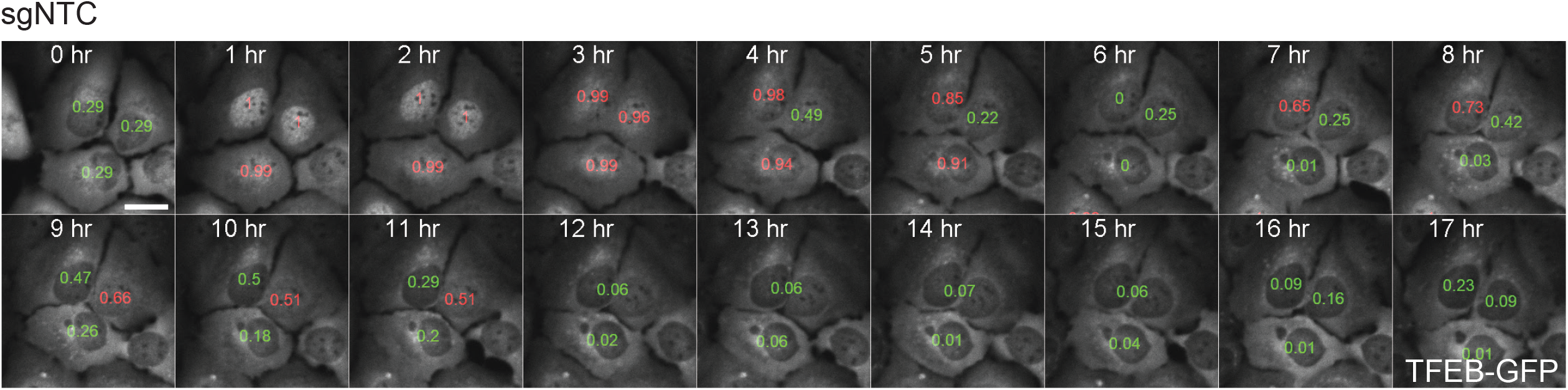
Live-cell images of TFEB-GFP U2OS cells under starvation conditions. TFEB-GFP-expressing U2OS cells were starved for 18 hours and images acquired every hour. low probability values are shown in green for cytosolic TFEB-GFP, high probability values (red) for nuclear TFEB-GFP. sgNTC, nontargeting sgRNA control, scale bar 5 μm

**Figure S5.**
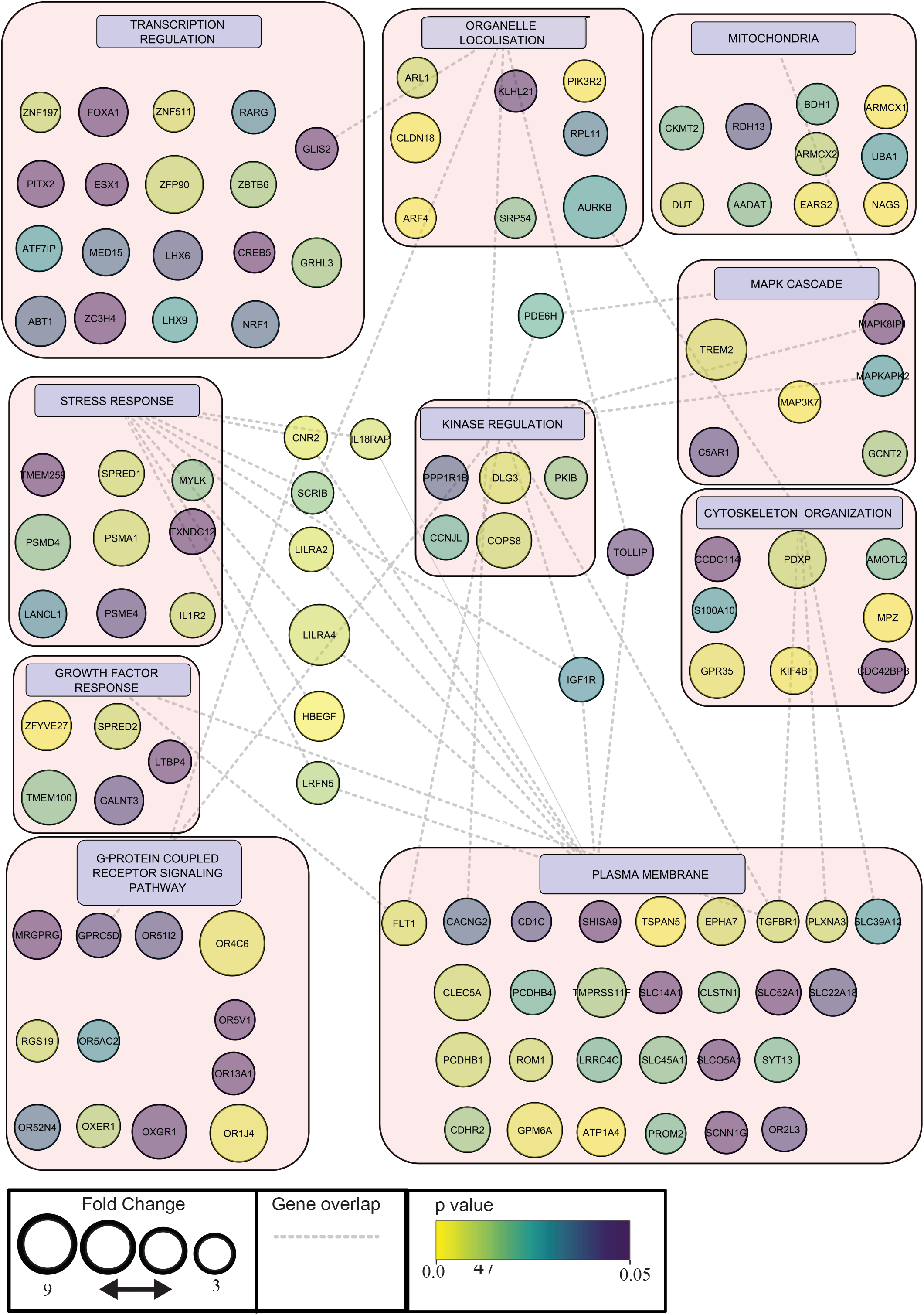
Gene-set clustering of 64 candidates enriched in TFEB-GFP translocation screen. Cytoscape analysis of enriched genes, the circle size represents fold-change enrichment and p-value is color-coded inside the circle. The dashed lines indicate cluster overlap.

Video 1 | **Example of AI-PS platform Parkin screen proof of principle**, Nikon NIS elements JOB module screen capture during GFP-Parkin (in Green) acquisition. Machine learning deployed for automatic detection of sgRNA targeting PINK-1 (in blue) according to cell phenotype (red circle on top of GFP-Parkin Image). The detected cell is photoactivated (in yellow-red).

Video 2 | **Live-cell images of TFEB-GFP U2OS cells under starvation conditions.** TFEB-GFP-expressing U2OS cells were starved for 18 hours and images acquired every hour. Single-cell CNN prediction scores are marked in red for nuclear TFEB and in green for cytosolic TFEB. Dynamic Bar chart indicates the cumulative distribution of TFEB translocation in the represented cell population.

Video 3 | **Example of AI-PS platform for TFEB screen**, Nikon NIS elements JOB module screen capture during TFEB-GFP (in green) acquisition. Top Image TFEB-GFP in green, red circle for the phenotype automatic detected cells. Bottom left corner, R based image segmentation, machine learning prediction and Mask generation. Bottom right, photoactivation in red (pa-mCh). Three examples are shown here.

Video 4 | **Live-cell images of TFEB-GFP U2OS cells expressing sgRNA targeting TGFBR1 under starvation conditions.** sgTGFBR1-TFEB-GFP-expressing U2OS cells were starved for 18 hours and images acquired every hour. Single cell CNN prediction scores are marked in red for nuclear TFEB and in green for cytosolic TFEB. Dynamic Bar chart indicates the cumulative distribution of TFEB translocation in the represented cell population.

Video 5 | **Live-cell images of TFEB-GFP U2OS cells expressing sgRNA targeting CREB5b under starvation conditions.** sgCREB5b-TFEB-GFP-expressing U2OS cells were starved for 18 hours and images acquired every hour. Single cell CNN prediction scores are marked in red for nuclear TFEB and green for cytosolic TFEB. Dynamic Bar chart indicates the cumulative distribution of TFEB translocation in the represented cell population.

Video 6 | **Live-cell images of TFEB-GFP U2OS cells expressing sgRNA targeting PPP1R1B under starvation conditions.** sgPPP1R1B-TFEB-GFP-expressing U2OS cells were starved for 18 hours and images acquired every hour. Single cell CNN prediction scores are marked in red for nuclear TFEB and in green for cytosolic TFEB. Dynamic Bar chart indicates the cumulative distribution of TFEB translocation in the represented cell population.

Video 7 | **Live-cell images of TFEB-GFP U2OS cells expressing sgRNA targeting mTOR under starvation conditions.** Sg-mTOR-TFEB-GFP-expressing U2OS cells were starved for 18 hours and images acquired every hour. Single cell CNN prediction scores are marked in red for nuclear TFEB and in green for cytosolic TFEB. Dynamic Bar chart indicates the cumulative distribution of TFEB translocation in the represented cell population.

Table S1. **Parkin translocation screen using an sgRNA library subpool targeting all kinases, phosphatases and the druggable genome.** The rows in the Parkin screen gene enrichment spreadsheet are the proteins selected by the screen. Log2 fold change and corresponded p-values calculated from the gene abundance analysis as described in Methods and related to Figure 3.

Table S2. **TFEB translocation whole genome screen.** The rows in the TFEB screen gene enrichment spreadsheet are the proteins selected by the screen. Log2 fold change and corresponded p-values calculated from the gene abundance analysis as described in Methods and related to Figure 5.

